# Inactivation of phosphodiesterase-4B gene in rat nucleus accumbens shell by CRISPR/Cas9 modulates the motivation to chronically self-administer nicotine

**DOI:** 10.1101/2023.03.07.531588

**Authors:** Burt M Sharp, Qin Jiang, Panjun Kim, Hao Chen

**Affiliations:** Department of Genetics, Genomics and Informatics, College of Medicine, University of Tennessee Health Science Center, Memphis, Tennessee; Department of Pharmacology, Addiction Science and Toxicology, College of Medicine, University of Tennessee Health Science Center, Memphis, Tennessee

## Abstract

Large scale human genome wide association studies (GWAS) have identified a growing pool of genes associated with cigarette smoking. One of the most prominent, phosphodiesterase-4B (PDE4B), has been associated with multiple smoking phenotypes. Although PDE4B modulates the half-life of neuronal cAMP, its precise role in smoking behaviors is unknown. To address this knowledge gap, we used a reverse translational approach. We inactivated *PDE4B* in bilateral medial nucleus accumbens shell (NAcs) neurons by injecting AAV containing a specific gRNA in female transgenic Cas9+ Long Evans rats. These rats then were given 23-hour chronic access to nicotine intravenous self-administration (IVSA) under a schedule of increasing fixed ratios (FR). With the increased effort required at FR7, nicotine SA (i.e. active presses and drug infusions) declined significantly in controls, whereas it was maintained in the mutagenized group. A progressive ratio (PR) study also showed significantly greater cumulative nicotine infusions in the mutant group. Hence, we hypothesized that enhanced PDE4B protein activity would reduce nicotine IVSA. A positive allosteric modulator,2-(3-(4-chloro-3-fluorophenyl)-5-ethyl-1H-1,2,4-triazol-1-yl)-N-(3,5-dichlorobenzyl)acetamide (MR-L2), was microinfused into NAcs bilaterally at FR3 or FR5; in both cohorts, MR-L2 acutely reduced nicotine IVSA. In summary, these studies show that the activity of PDE4B regulates the capacity of NAcs to maintain nicotine IVSA in face of the cost of increasing work. This finding and the results of the PR study indicate that PDE4B affects the motivation to obtain nicotine. These reverse translational studies in rats provide insight into the motivational effects of NAcs PDE4B that advance our understanding of the smoking behaviors mapped in human GWAS.

## Introduction

Human genome wide association studies (GWAS) have identified a growing pool of genes associated with cigarette smoking (1,2). A study of 1.2 million individuals in the British Biobank identified phosphodiesterase-4B (PDE4B) as one of only three genes (2) independently associated with multiple smoking phenotypes including initiation of regular smoking, age of regular smoking, amount smoked (cigarettes/day), and smoking cessation (1,2). Recently, meta-analysis of a GWAS involving 60 ancestrally diverse cohorts, comprised of 3.4 million individuals, found seven distinct single nucleotide polymorphisms (SNPs) in an intron of phosphodiesterase-4B (PDE4B) - each associated with smoking initiation across ancestral groups (2). As genes associated with tobacco smoking are confirmed by human GWAS, understanding the functional significance of specific genes has become increasingly important in the rational selection of novel therapeutic targets to improve the success of smoking cessation (2,3). Reverse translational modeling of human GWAS findings in established animal models of addiction is a potent tool for understanding the behavioral function of specific genes (4).

PDE4B belongs to a multi-gene family of PDE isoenzymes that regulate cyclic nucleotide homeostasis in multiple intracellular compartments (5). PDE4B is one of four PDE4 isoenzymes (A,B,C,D) encoded by four genes that specifically dephosphorylate cyclic AMP (cAMP), selectively limiting the half-life of cAMP-dependent signaling by signaling complexes involved in specific cellular processes (6,7). Therefore, the PDE4B expressed in nucleus accumbens (NAc) (8), the brain region mediating the motivation to take drugs including nicotine (9), is most likely a critical regulator of signaling pathways controlling the activation of median spiny neurons (MSNs) in NAc.

The neural circuitry of NAc, largely comprised of MSNs, functions as the brain interface mediating the motivation for goal directed behaviors by integrating information from cortical and limbic inputs with outputs to brain regions involved in goal-directed behaviors (e.g., ventral pallidum). By regulating the half-life of cAMP associated with signaling complexes in MSNs, PDE4B affects the activity of key downstream signaling molecules including protein kinase A (PKA), dopamine and cAMP-regulated phosphoprotein (DARPP-32), and cAMP-response element binding protein (CREB) (10,11). These signaling molecules modulate the excitability of MSNs (12–14). Signaling in this pathway (i.e., cAMP/PKA/DARPP-32/CREB) is activated by dopamine receptors (D1R), in response to dopamine release induced by nicotine and other drugs of abuse, and inhibited by D2R; these dopamine receptor (DaR) subtypes are differentially expressed by two types of MSNs (15–17).

The effects of modulating PDE4B activity on operant intravenous self-administration (IVSA) of nicotine are unknown. Based on human GWAS that have demonstrated the association of PDE4B with human smoking behavior (1,2), we addressed the role of PDE4B in operant nicotine IVSA by using a well-established rat model of virtually unlimited chronic access to the drug (18,19). We sought to disrupt PDE4B gene expression in medial NAc shell (NAcs) neurons, by *in vivo* genome editing using virally delivered gRNA targeting all transcripts of PDE4B in transgenic rats containing CRISPR/Cas9 (20). To accomplish this and to selectively target neurons controlling motivation that project to ventral pallidum, rats received bilateral medial NAcs microinjections of adeno-associated virus (AAV) containing the PDE4B gRNA. In retrograde transport studies, we had previously shown that medial NAcs neurons project to ventral pallidum (21). Additional experiments were designed to enhance the activity of PDE4B protein in NAc by administering a positive allosteric modulator (PAM) specific for PDE4 long isoenzymes, which contain two upstream conserved regulatory regions (i.e., UCR1 and UCR2) (7,22). This PAM, 2-(3-(4-chloro-3-fluorophenyl)-5-ethyl-1H-1,2,4-triazol-1-yl)-N-(3,5-dichlorobenzyl)-aceta-mide (MR-L2), targets UCR1 and has been shown to reduce cellular cAMP levels and PKA signaling (7,22). We found that inactivating the PDE4B gene in NAcs neurons by CRISPR/Cas9-induced mutation increased the motivation to obtain nicotine as the FR requirement increased (i.e., number of operant lever presses required to obtain a single i.v. infusion of nicotine), whereas administration of MR-L2 had the opposite effect - decreasing the motivation to obtain nicotine.

## Methods

### Materials, Animals and Breeding

Heterozygous loxP-STOP-loxP Cas9+ female outbred Long Evans rats, originally developed by Brandon K. Harvey Ph.D. at NIH (20), and provided by Dr. Aron Geurts at Medical College of Wisconsin, were used in all experiments. We maintained a breeding colony using wildtype Long Evans rats from Charles River, and genotyped (Transnetyx, Memphis, TN) to identify Cas9+ females. 2-(3-(4-chloro-3-fluorophenyl)-5-ethyl-1H-1,2,4-triazol-1-yl)-N-(3,5-di-chlorobenzyl)acetamide (MR-L2) was from Targetmol Chemicals, Wellesley Hills, MA (CAS#2374703-19-0). An MR-L2 stock solution (5.0 mM) in 100% DMSO was diluted with 20% Sulfobutylether-β-Cyclodextrin (SBE-β-CD; CAS #182410-00-0; Targetmol Chemicals Inc.) in sterile saline.

### Identification of an effective gRNA complementary to rat PDE4

In consultation with Applied Biological Materials Inc., Richmond, BC, Canada, we designed three gRNAs to rat PDE4B, using their proprietary software, and generated viral vectors (10^12^GC/ml) with the following structure: pAAV-hSyn-Kozak-iCre-P2A-mClover3-U6-gRNA. Each was tested by microinfusion (400nl) into prefrontal cortex. We initially tested 40nl (4×10^4^ genome copy of AAV, n=2), and then 400nl injections (4×10^5^ genome copy of AAV, n=4). We sampled from 4-5 coronal tissue sections (100 μm), based on fluorescence intensity of mClover3 in sentinel sections; genomic DNA extraction from a region surrounding the injection site; and PCR amplification (5’ primer: gacagcaaaagtcacatgcag; 3’: gaacaaatggggccttaaca). Amplification of gDNA was performed via Hi-Plex approach optimized by Floodlight Genomics (Knoxville, TN), followed by multiplex sequencing using a single end 150 kit on an Illumina HiSeq X (San Diego, CA). Data was analyzed by CRISPRpic (23) to identify and classify mutations. Only one gRNA, TGCATGTGAGGGGCCGATTA, effectively mutated the PDE4B DNA. This gRNA is on chr5:122128512-122128531 (Rnor6.0) and chr5:117359381-117359400 (mRatBN7.2).

### Stereotaxic surgery

The same procedure was used for: (i) *in vivo* CRISPR/Cas9 mutagenesis of neuronal PDE4B genomic DNA in nucleus accumbens shell (NAcs); (ii) implantation of NAcs guide cannulae to deliver MR-L2. To avoid the considerable day-to-day variation in nicotine SA observed in pilot studies of male LE rats, females were used throughout these studies.

Female Cas9+ LE rats, 180-200g b.wt., were anesthetized with 2% isoflurane and microinfused with AAV or control constructs (AAV-hU6-GLuc219-gRNA-EF1α -EGFP-KASH; 3.68E+12 GC/ml; from Dr. Brandon Harvey) (20) targeted bilaterally to medial NAcs (AP 1.8; ML 1.4 or -1.4; DV 6.7; 5 degree). For administration of MR-L2 during nicotine IVSA, rats were implanted with guide cannulae in medial NAcs bilaterally and affixed to the skull with screws and dental cement. Ten days later, while anesthetized, a jugular vein cannula was implanted and tunneled subcutaneously to a button affixed to an incision on the back; a spring-loaded tether was attached to deliver nicotine. All procedures and protocols were approved by the IACUC of UTHSC.

### Access to virtually unlimited operant nicotine SA

Rats were individually housed in operant chambers and given access to nicotine SA (23h/day, 7 days/week) without food deprivation or training (18,19). Operant chambers contained two horizontal levers; during IVSA sessions, a green cue light above each lever signaled nicotine was available. Lever presses were recorded and syringe pumps were controlled by computers, using GS3 software (Coulbourn Instruments, Allentown, PA). Nicotine IVSA began on initiation of the dark cycle (10 AM/12 hours). Pressing the active lever elicited a nicotine injection (0.03 mg/kg per 50μl/0.81s, free base, pH 7.2–7.4). To prevent an overdose, green cue lights were extinguished and nicotine was unavailable for 7s after each nicotine injection. Pressing the inactive lever had no programmed consequence. The final hour of the 12 h lights-on cycle (9:00–10:00 AM) was reserved for animal husbandry. Catheter patency was checked with brevital injection (0.2 ml; JHP pharmaceuticals, Rochester, MI). Rats with occluded catheters were excluded from study.

### Experiment 1: Increasing fixed ratio (FR) required to obtain nicotine in rats with bilateral PDE4B mutations in medial NAcs vs. controls

Stable nicotine intake in daily 23 hour sessions was achieved within 10 days of initiating nicotine IVSA at FR1 (1 active lever press triggered 1 nicotine infusion); thereafter, the FR requirement was advanced as follows: FR2, 5 days; FR3, 5 days; FR5, 7 days; FR7, 7 days. Thereafter, cardiac perfusion under lethal isoflurane was performed with 4% paraformaldehyde, then brains were removed and dehydrated in 30% sucrose for 2 days. Coronal cryostat sections (100μm) were obtained from all rats for fluorescence microscopy (excitation, 505nm; emission, 515nm); sections were evaluated from all rats.

### Experiment 2: Effects of bilateral NAcs infusions of MR-L2 vs. vehicle control during operant nicotine IVSA at increasing FR requirements

The FR schedule was: FR1, 10 days; FR2, 5 days, FR3, 5 days, FR5, 9 days. MR-L2 or vehicle was administered on day 4 of FR3 in one group and on day 8 of FR5 in a separate group. During the final hour of lights-on, microinfusion needles were inserted into guide cannulae while rats were briefly anesthetized with 2% isoflurane; anesthesia ceased, MR-L2 (300 nl; 2×10^-4^ M; 1:20, DMSO in 20% SBE-β-CD) or vehicle (1:20, DMSO in 20% SBE-β-CD) was microinfused (25 nl/min over 12 min) into NAcs, and needles were removed 5 min later. Rats returned to home operant chambers 20 min later, the dark phase of the light cycle began, and nicotine was available. Following the experiment, brains were removed from rats treated with lethal isoflurane + thoracotomy. Brains were flash frozen, maintained at −20℃, and cryosectioned (50 μm) to locate guide cannulae.

### Data Analysis and Statistics

Active lever presses and nicotine infusions at FR7 and during a PR (progressive ratio) study were compared in AAV experimental and control groups. For studies of MR-L2, similar parameters were compared during FR3 or FR5 in independent experiments. Cumulative active presses and infusions were divided into consecutive 3-hour bins on the day that MR-L2 or vehicle was injected in each experiment. These data were expressed as a percentage of the mean value for the corresponding time bins recorded during the 3 preceding days.

Data were analyzed by repeated measures 2-way ANOVA, using R package, with either day of lever press or nicotine infusion designated as a within-subject factors and treatment designated as a between-subject factor. Tukey HSD test was used for *post hoc* testing. PR data and time binned data were compared between treatment groups by t-test; P values were Bonferroni-adjusted for multiple comparisons. Data were expressed as mean ± SEM. Statistical significance was assigned at *p* < 0.05.

## Results

Using the CRISPR/Cas9 technology targeted to neurons by neuron-specific expression of iCre and, therefore, Cas9, a gRNA specific for rat PDE4B exon 9, common to all PDE4B transcripts, generated several types of edits (Figure 1) detected in heterogeneous tissue dissected from the medial NAcs after AAV injection. We selectively introduced loss of function edits into the PDE4B gene of NAcs neurons in Loxp-STOP-Loxp-CRISPR/Cas9 knockin rats (20). Neuronal specificity was achieved by injecting an AAV vector containing an hSyn promoter, which drives the expression of iCre, and a gRNA targeting exon 9 of PDE4B, present only in PDE4B transcripts (i.e, pAAV8-hSyn-Kozak-iCre-P2A-mClover3-U6-gRNA). DNA was extracted from heterogeneous tissue dissected from the medial NAcs 10 days after AAV injection. The genomic sequence flanking the gRNA target site was amplified using PCR and sequenced on an Illumina NovaSeq instrument. This tissue is approximately two thirds glial and one third neuronal (24). However, only a relatively small subset of cells from the neurons infected by the AAV would yield mutant sequences. Considering the diffusion volume of the AAV infusate and the injection site, we estimated that only 25-35% of the neurons in the sampled tissue punch could be transfected by the AAV.

**Figure 1.**
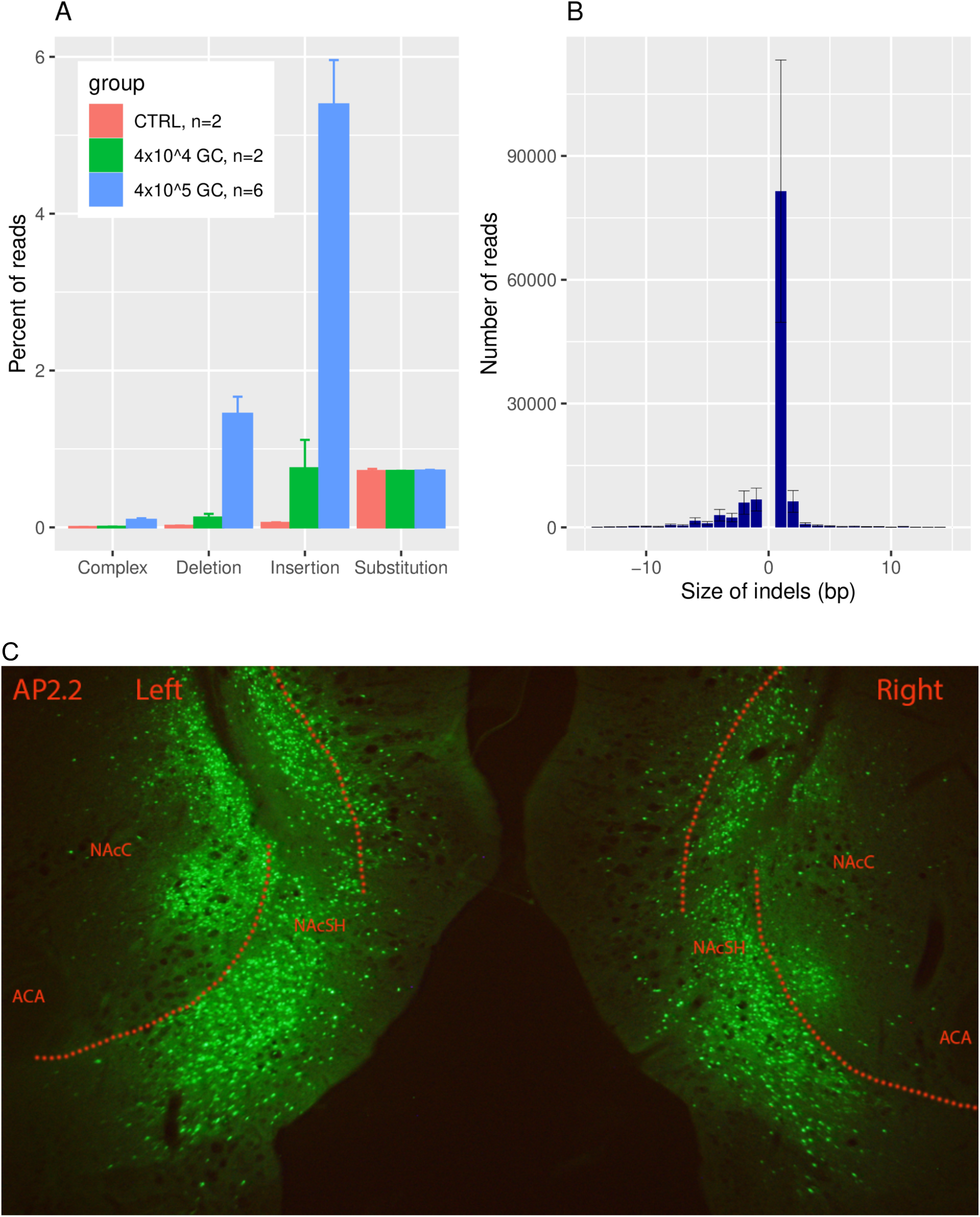
Detection of genome editing and photomicrographs of mClover fluorescence in NAcs. To characterize the genome editing, prefrontal cortex (PFC) was microinjected *in vivo* with pAAV8-hSyn-Kozak-iCre-P2A-mClover3-U6-gRNA or control construct and tissue was obtained 10 days later (panels A, B). Expression of mClover3 (panel C) was detected in NAcs 45 days post-injection of pAAV8 construct, following completion of a trial of nicotine SA through FR7. A. The gRNA targeting exon 9 of PDE4B induced mutations in PFC genomic DNA in a dose-dependent manner. The percent of reads that contain mutations is in line with expectations for a heterogeneous tissue. Complex mutations contain both insertion and deletion. GC: Genome copy. B. The size of indels range from −81 to 79 bp. The vast majority (95+/-0.5%) of these mutations will produce frame-shifts, leading to loss of protein function. C. mClover fluorescence demonstrates strong expression of viral vector in NAcs neurons.

Since one third of the tissue was neuronal and 25-35% of all neurons could be transfected, approximately 8-12% of all cells were actually targeted for editing. In line with this expectation, approximately 6.9+/-1.9 percent of the sequenced reads contained mutations in rats that received the PDE4B gRNA injection (Figure 1A-a). By far, the most frequent type of mutation is 1 bp insertion, followed by 1 bp deletion (Figure 1A-b). In total, mutations range in size from −81 to 79 bp. Each of these types of mutation, i.e., insertions, deletions and complex changes in PDE4B, were approximately 95, 75 and 20-fold, respectively, more common in tissue from NAcs treated *in vivo* with PDE4B-gRNA compared to control (AAV-hU6-GLuc219-gRNA-EF1α-EGFP-KASH) (20). The fold-change depended on the dose of AAV injected. Insertions, for example, were reduced from 95-fold to 13-fold greater than control by decreasing the dose 10-fold. The vast majority (95+/-0.5%) of these mutations are known to produce frameshifts, which result in the loss of protein function.

A representative photomicrograph of mClover fluorescence from one rat brain at AP 2.2 is shown in Figure 1B. Abundant fluorescence, similar to that detected in all rats 45 days after injection of the AAV PDE4B-gRNA construct and completion of a trial of nicotine SA through FR7, is predominantly in nuclei within the NAc shell of this representative sample. Hence, Cas9-dependent mutagenesis, targeted to PDE4B by the PDE4B-gRNA, is localized to neurons largely in the medial NAc shell.

### Experiment 1

Figure 2 shows active lever presses (top 2 panels) and nicotine infusions by day as FR increased from FR1 to FR7. By the final 2-3 days at FR1-5, presses and infusions were similar in the experimental (EXP) and control (CNT) groups. However, at FR7, these parameters diverged between the two treatment groups: both presses and infusions increased from the first to the final day at FR7 in the EXP group, whereas both parameters decreased over this time interval in the CNT group. Comparing the final 3 days of active lever presses at FR7 to FR5, in the EXP group FR7 was greater, whereas it was less in the CNT group. Nicotine injections were similar during the final 3 days of FR7 vs. FR5 in the EXP group, whereas injections declined in the CNT group.

**Figure 2.**
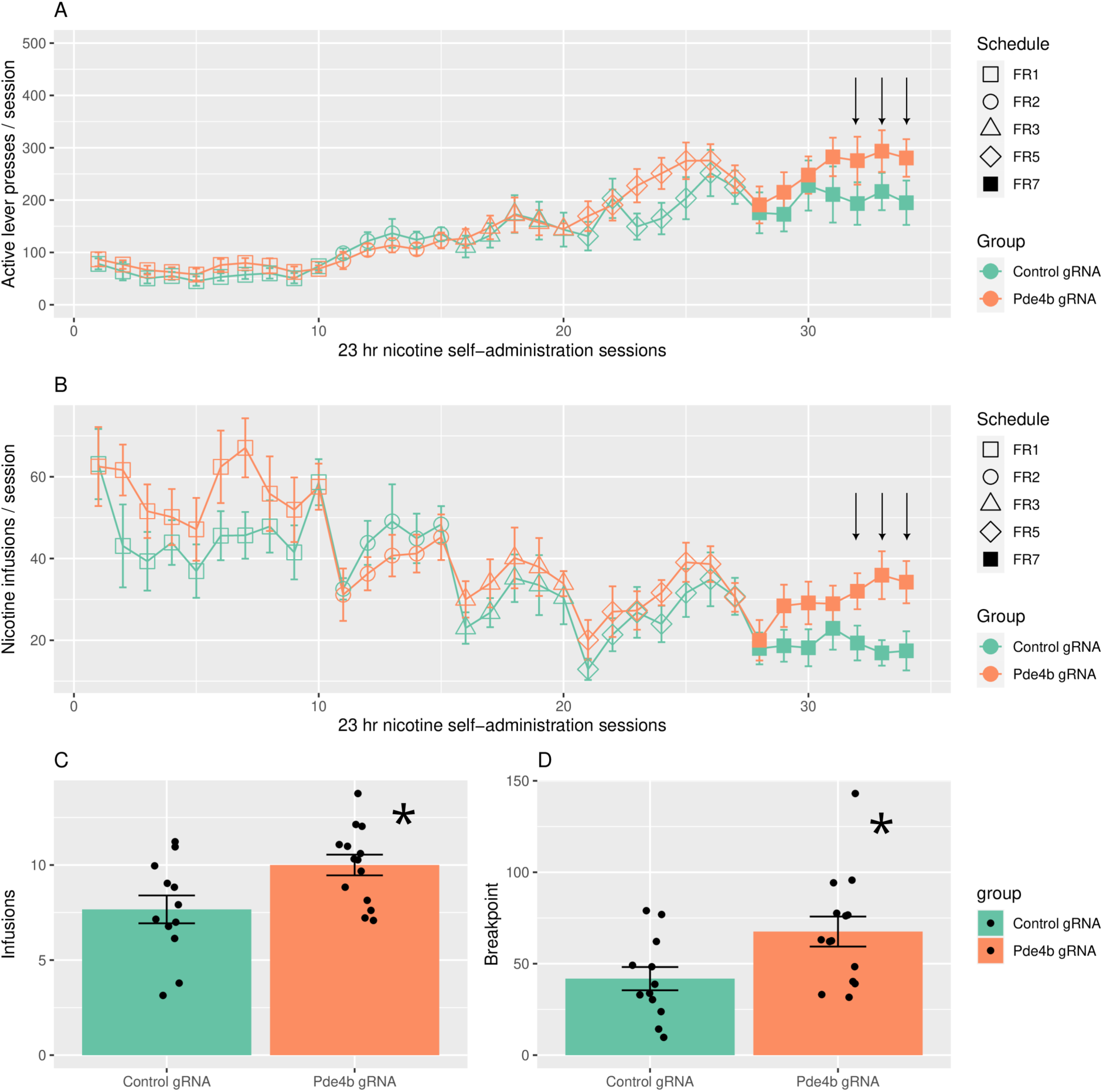
Operant nicotine SA under an increasing fixed ratio (FR) schedule, progressive ratio (PR) test, and NAcs locations of injection needle tips in rats treated with an AAV-PDE4B-gRNA (EXP) or control (CNT) constructs. Rats had access to nicotine SA everyday for 23 hours/day in operant chambers equipped with active and inactive levers. Daily active lever presses (A) and nicotine injections (B) are shown according to FR schedule for both treatment groups. The average daily active press elicited by each rat during the final 3 days of FR7 (marked by arrows): EXP = 403.1 ± 68.9 (mean ± SE; n=14) vs. CNT = 211.3 ± 36.5 (n=12) (P=0.028); nicotine infusions, EXP = 33.9 ± 4.6 vs. CNT = 17.9±3.7 (P=0.0134). Analysis of all 7-daily FR7 responses for EXP vs CNT (repeated measures 2-way ANOVA): showed a significant treatment effect on injection (P=0.046), but not on active presses (P=0.08). In the PR test, conducted a day after completion of nicotine SA at FR7, the cumulative infusions (C) obtained by the two treatment groups were: EXP = 10.0 ± 0.5 (mean ± SE; n=14) vs. CNT = 7.7 ± 0.7 (n=12) (*, P<0.02; unpaired T-test, 2-tailed). In addition, the breakpoint (D) achieved by the EXP group (67.6 ± 8.2 active presses for PDE4B gRNA) was significantly greater than CNT (41.8 ± 6.4 active presses; *, P<0.02).

Taking the average daily active press elicited by each rat during the final 3 days of FR7, we found: EXP = 403.1 ± 68.9 (mean ± SE; n=14) vs. CNT = 211.3 ± 36.5 (n=12) (P=0.028); nicotine infusions, EXP = 33.9 ± 4.6 vs. CNT = 17.9±3.7 (P=0.0134). Comparing all 7 daily FR7 responses for EXP vs CNT (repeated measures 2-way ANOVA) showed a significant treatment effect on injection (P=0.046), but not on active presses (P=0.081). Lastly, a progressive ratio (PR) test was conducted the day after completion of nicotine IVSA at FR7. Figure 2 (lower panels: a, b) shows the cumulative infusions and breakpoints achieved in the two treatment groups: infusions, EXP = 10.0 ± 0.5 vs. CNT = 7.7 ± 0.7 (P=0.018); breakpoints, EXP = 67.6 ± 8.2 active presses vs CNT = 41.8 ± 6.4 (P=0.02). In summary, region-specific mutagenesis of PDE4B in medial NAcs sustained lever pressing to self-administer i.v. nicotine despite the high FR requirement (i.e. FR7), whereas active presses and nicotine infusions declined in the CNT group. This difference is reinforced by the significant difference in infusions and breakpoints achieved during the PR test - an index of motivation to obtain nicotine.

### Experiment 2

Rats with bilateral NAcs guide cannula implants self-administered nicotine under a schedule of increasing FR. Figure 3(C) is a schematic showing the NAcs positions of the injection needle tips used to administer MR-L2 (2×10^-4^ M, 300nl/side) on day 6 of the 7-day FR3 schedule. Almost all injection tips were tightly localized to medial NAcs.

**Figure 3.**
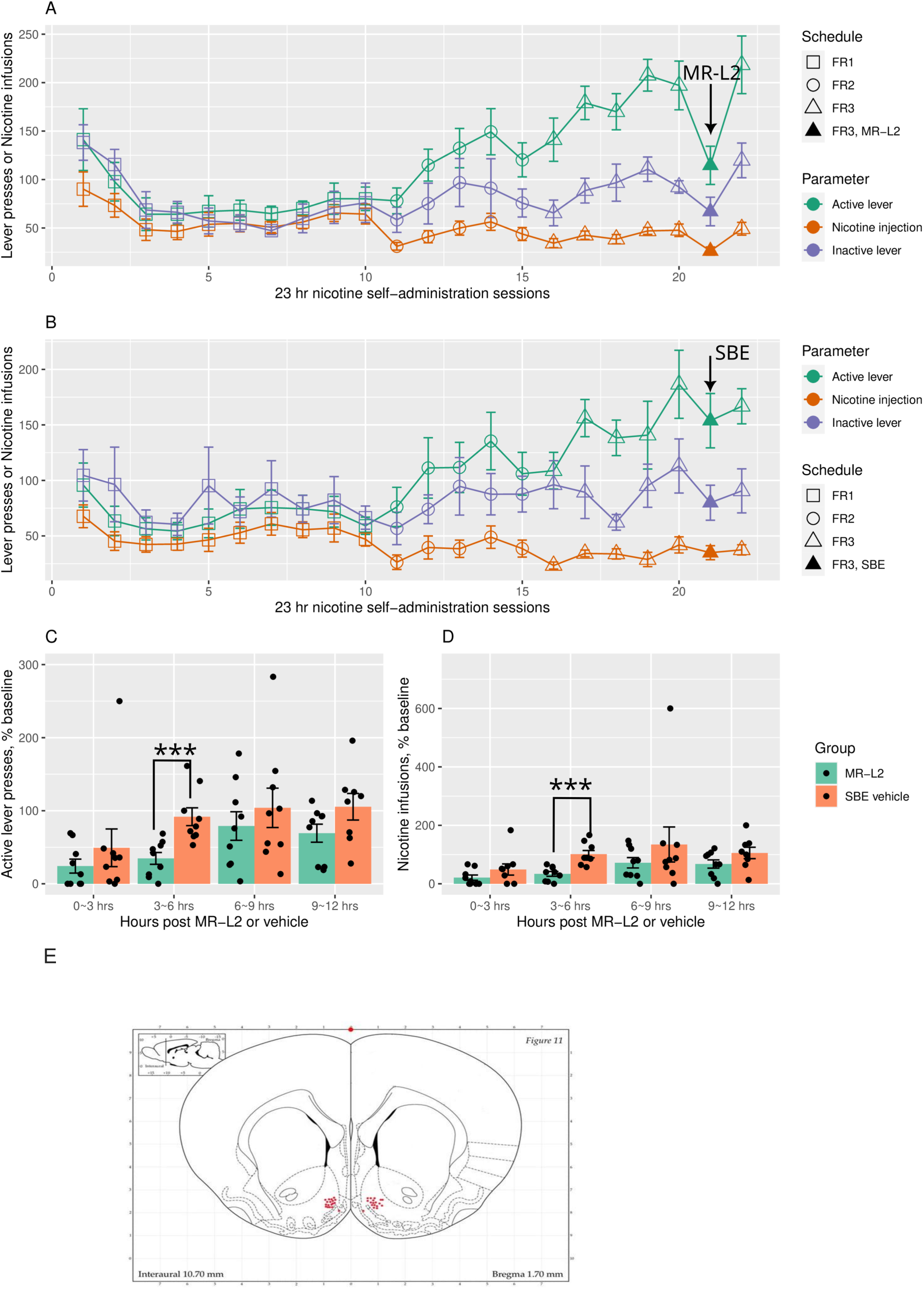
Operant nicotine SA under an increasing fixed ratio (FR) schedule in rats treated with MR-L2 (EXP) or vehicle (CNT) at FR3. Active and inactive presses and infusions are shown for EXP (A. MR-L2 2×10^-4^ M, 300nl/side) and CNT (B. DMSO-SBE) groups. Comparing days 3-6, MR-L2 significantly reduced active presses on day 6 in EXP, but not in CNT: effect of treatment, F_1,16_ =0.49, P=0.5; day, F_1,16_ =10.35, P=0.0054; treatment x day, F_1,16_ =9.60, P=0.0069 (repeated measures 2-way ANOVA). *Post-hoc* analysis showed that active presses on days 3-5 were greater than day 6 in the EXP (P=0.029, Tukey), but not CNT (P=0.99) group. Nicotine infusions were significantly reduced by MR-L2 on day 6: treatment, F_1,16_=0.69, P=0.42; day, F_1,16_ =9.28, P=0.0077; treatment x day, F_1,16_ =9.44, P=0.0073; infusions on days 3-5 > day 6 in EXP (P=0.033; Tukey), but not CNT (P=0.99). Inactive presses were not different by treatment group (F_1,16_ =0.07, P=0.795) nor was an effect of day detected (F_1,16_ =3.79, P= 0.070). Total active presses elicited or infusions were binned in time intervals 0-3, 3-6, 6-9, and 9-12 hours after administration of MR-L2 into NAcs bilaterally. These data were normalized by the average value for the corresponding time bins recorded on days 3-5. Significant effects of MR-L2 (EXP) vs. vehicle (CNT) on active presses (C) and infusions (D) occurred between 3-6 hours after dosing: treatment at 3-6 hours post MR-L2, P=0.0013 and 0.0003, respectively; all other time bins, P>0.05 (P values adjusted for multiple testing). A schematic (E) showing the NAcs locations of all injection needle tips used to administer MR-L2 to rats on day 6 of the 7-day FR3 schedule. The brain section was adapted from the rat brain atlas of Paxinos and Watson (46).

Figure 3(A) shows the effect of MR-L2 on active presses and infusions at FR3 in EXP (top panel) and CNT (i.e., received DMSO-SBE) groups. MR-L2 significantly reduced active lever presses on day 6 in EXP, but not in CNT: effect of treatment, F_1,16_ =0.49, P=0.5; day, F_1,16_ =10.35, P=0.0054; treatment x day, F_1,16_ =9.60, P=0.0069 (repeated measures 2-way ANOVA). *Post-hoc* analysis showed that active presses on days 3-5 were greater than day 6 in the EXP (P=0.029, TukeyHSD), but not CNT (P=0.99) group. We found similar differences for nicotine infusions between the two treatment groups: treatment, F_1,16_ =0.69, P=0.42; day, F_1,16_ =9.28, P=0.0077; treatment x day, F_1,16_ =9.44, P=0.0073; infusions on days 3-5 > day 6 in EXP (P=0.033; TukeyHSD), but not CNT (P=0.99). Inactive presses were not different by treatment group (F_1,16_ =0.07, P=0.795) nor was an effect of day detected (F_1,16_ =3.79, P= 0.070).

We further analyzed the effects of MR-L2 by generating time bins containing all active presses elicited or infusions obtained within time intervals 0-3, 3-6, 6-9, and 9-12 hours after administration of MR-L2. These data were normalized by the average value for each time bin recorded on days 3-5. Figure 3(B) shows that the greatest effects of MR-L2 (EXP) vs. vehicle (CNT) on active presses and infusions occurred between 3-6 hours after dosing: effects of treatment at 3-6 hours post dose, P=0.0048 and 0.0012, respectively; all other time bins, P>0.05 (Bonferroni-adjusted P values).

Figure 4(C) is a schematic showing the NAcs positions of the injection needle tips used to administer MR-L2 (2×10^-4^ M, 300nl/side) on day 8 of the 9-day FR5 schedule. The effects of MR-L2 on active presses and nicotine infusions at FR5 in EXP and CNT (i.e., DMSO-SBE) groups are shown in Figure 4(A). MR-L2 significantly reduced active presses on day 8 in EXP, but not in CNT: effect of treatment, F_1,21_=0.085, P=0.74; day, F_1,21_=20.63, P=0.00018; treatment x day, F_1,21_=3.96, P=0.059. *Post-hoc* analysis showed that active presses on days 5-7 were greater than day 8 in the EXP (P=0.0071, Tukey), but not CNT (P=0.43) group. We found similar differences for nicotine infusions between the two treatment groups: treatment, F_1,21_=0.036, P=0.56; day, F_1,21_=18.12, P=0.00035; treatment x day, F_1,21_=8.35, P=0.0088; infusions on days 5-7 > day 8 in EXP (P=0.0029; Tukey), but not CNT (P=0.65). Inactive presses were not different by treatment group (F_1,21_=0.061, P=0.45), and an effect of day, but not treatment x day was detected (F_1,21_ = 16.64, P=0.0005; F_1,21_ = 0.067, P=0.80, respectively). However, *post-hoc* analyses showed that inactive presses on days 5-7 were not different from day 8 in the EXP (P=0.10, Tukey) and CNT (P=0.29) groups.

**Figure 4.**
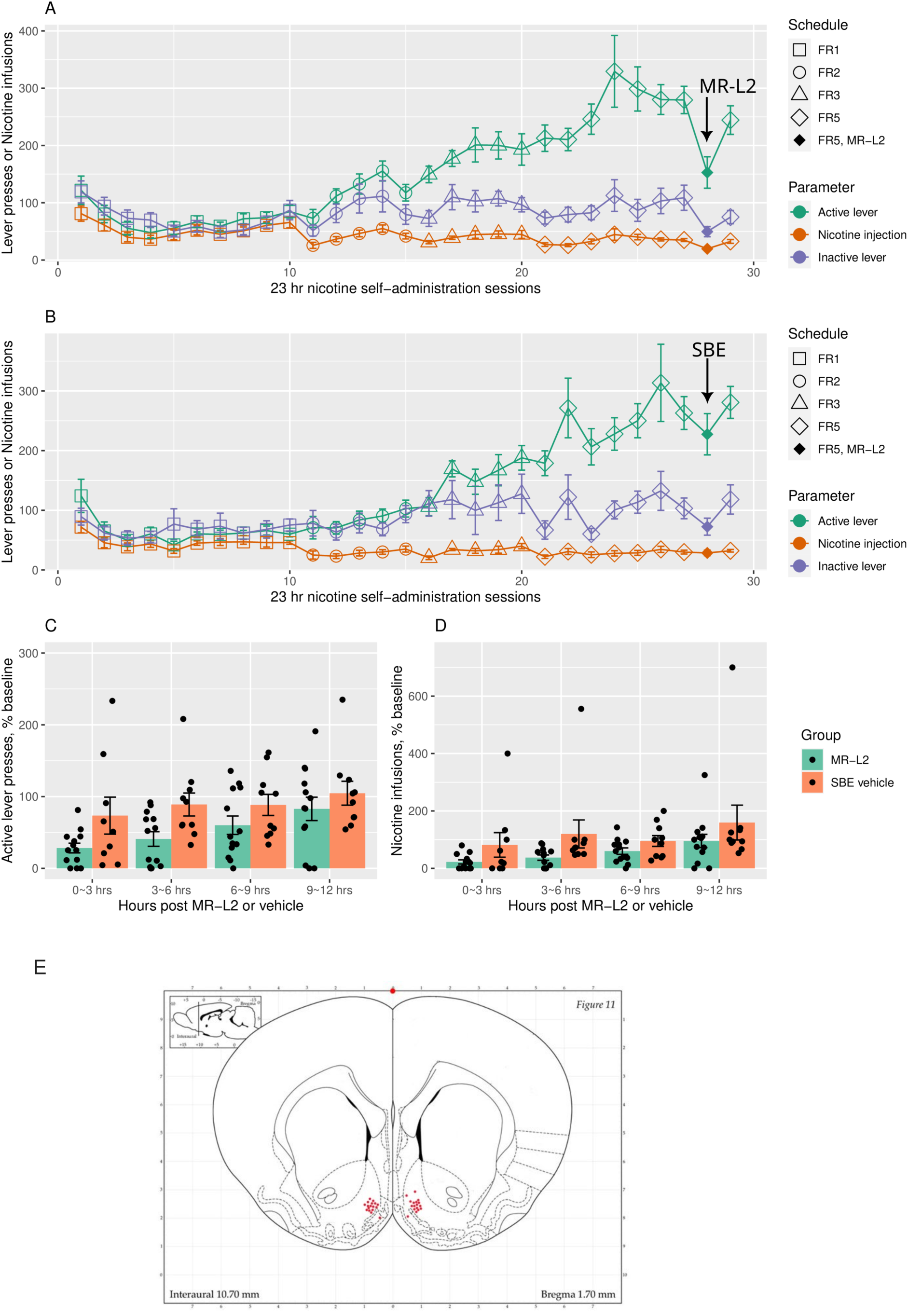
Operant nicotine SA under an increasing fixed ratio (FR) schedule in rats treated on day 8 with MR-L2 (EXP) or vehicle (CNT) at FR5. Active and inactive presses and infusions are shown for EXP (A. MR-L2 2×10^-4^ M, 300nl/side) and CNT (B. DMSO-SBE) groups. Comparing days 5-8, MR-L2 significantly reduced active presses on day 8 in EXP, but not in CNT: treatment, F_1,21_ =0.085, P=0.74; day, F1,21=20.63, P=0.00018; treatment x day, F_1,21_=3.96, P=0.059 (repeated measures 2-way ANOVA). Nicotine infusions were significantly reduced by MR-L2 on day 8: treatment, F1,21=0.036, P=0.56; day, F_1,21_=18.12, P=0.00035; treatment x day, F_1,21_=8.35, P=0.0088. Post-hoc analysis showed: active presses on days 5-7 > day 8 in the EXP (P=0.0071, Tukey), but not CNT (P=0.43) group; infusions on days 5-7 > day 8 in EXP (P=0.0029; Tukey), but not CNT (P=0.65). Inactive presses were not different by treatment group (F_1,21_=0.061, P=0.45), and an effect of day, but not treatment x day was detected (F_1,21_ = 16.64, P=0.0005; F_1,21_ = 0.067, P=0.80, respectively). However, *post-hoc* analyses showed that inactive presses on days 5-7 were not different from day 8 in the EXP (P=0.10, Tukey) and CNT (P=0.29) groups. Total active presses elicited or infusions were binned in time intervals 0-3, 3-6, 6-9, and 9-12 hours after administration of MR-L2 into NAcs bilaterally. These data were normalized by the average value for the corresponding time bins recorded on days 5-7. A significant effect of treatment group was found for both active presses (C) and infusions (D): F_1,21_=7.06, P=0.015 and F_1,21_=6.34, P=0.02, respectively (repeated measures 2-way ANOVA); neither bin time nor treatment x time were significant (P > 0.05), nor were *post-hoc* tests. A schematic (E) showing the NAcs locations of all injection needle tips used to administer MR-L2 to rats on day 8 of the 9-day FR5 schedule.

Figure 4(B) shows the time bins for active presses and nicotine infusions obtained after administration of MR-L2 at FR5. A significant effect of treatment group was found for both presses and infusions: F=5.37, P=0.030 and F=4.84, P=0.038, respectively (repeated measures 2-way ANOVA); neither bin time nor treatment x time were significant (P > 0.05). *Post-hoc* tests were not significant.

## Discussion

Although human GWAS have implicated PDE4B in multiple stages of smoking behavior (1,2), the mechanism(s) underlying these effects are unknown. We hypothesized that reducing the function of PDE4B in NAcs neurons by CRISPR/Cas9 gene inactivation and enhancing the catabolic activity of the protein using MR-L2, a positive allosteric modulator, would provide insight into the role of PDE4B in nicotine SA under a schedule of increasing FR. Indeed, selective modulation of PDE4B by administration of the gRNA or MR-L2 into medial NAcs demonstrated significant and opposite effects on the maintenance of nicotine IVSA in face of increasing FR requirements. Mutagenesis of PDE4B by selective deletions and insertions in an exon specific for PDE4B maintained nicotine IVSA at high FR (i.e., FR7) compared to the declining IVSA observed in controls. Conversely, positive allosteric modulation of PDE4B rapidly reduced nicotine SA at FR3 and FR5; these FRs were otherwise well tolerated in controls.

These observations were made in rats that received microinfusions of an AAV construct into NAcs bilaterally - a construct designed to specifically edit PDE4B only in neurons. We tested our hypothesis in the NAcs because of its critical role in operant nicotine SA (9,25) and the relatively high concentration of PDE4B identified in this region (8). NAc is the brain interface that integrates mesocorticolimbic inputs, which transmit cognitive, emotive and reward-related information to NAc MSN. Monosynaptic inputs from regions such as frontal cortex, hippocampus and basolateral/basomedial amygdala (26), activate MSNs via glutamatergic projections. Additionally, nicotine-induced dopamine release from the ventral tegmental area to MSN modulates the activation of MSN (27). The NAc circuitry generates MSN outputs to regions such as ventral pallidum, which engage the brain circuitry involved in goal directed behaviors (28,29).

The high behavioral costs necessary to obtain rewards in certain operant paradigms are surmounted by increasing NAc neuronal activity (30). In high-cost trials, subsets of MSN fire prior to the initiation of behavioral responding and remain active until the reward is obtained, whereas the duration of responding is shorter when the cost is less (31). The costs of these predictably large responses are encoded by differences in phasic dopamine signaling to MSN (32). Dynamic changes in moment-to-moment dopamine release are critical to these cost-related decisions (33,34) involved in motivated behavior to obtain conditioned rewards. Based on the results of PDE4B mutagenesis, we hypothesize that the high cost of obtaining nicotine reward at FR7 is not met by increased MSN activity driven by enhanced dopamine release unless dopamine signaling is further amplified by molecular changes in the neuronal landscape, such as those induced by reducing the function of PDE4B in neurons.

PDE4B interacts with compartmentalized signaling complexes and modulates dopamine signaling by regulating the half-life of cAMP synthesized following D1R activation (5,34). Additionally, glutamatergic signaling to MSN AMPA receptors is modulated by D1R/cAMP/PKA (35–37). The duration and level of cAMP signaling determines the activity of downstream signaling moieties such as PKA, DARPP-32, PP1a and CREB (10,11), which are involved in regulating the excitation of MSN (12,14) and in regulation of the transcriptional landscape (38–40). Hence, PDE4B sits at a critical juncture - impacting both the acute and chronic activation of MSN.

The acute effects of modulating PDE4B activity and, thereby, MSN excitation are exemplified by the experiments with MR-L2. We observed the effects of this compound in rats self-administering nicotine at FR3 by 3 hours after infusion into NAcs (the administration protocol, which tended to reduce nicotine SA just after delivery of the drug, obviated detecting the effect of MR-L2 earlier), and the effects waned by 6 hours. Since positive allosteric modulation of PDE4B attenuates cAMP signaling, we would expect it to reduce nicotine and dopamine-dependent excitation of MSN (22). Indeed, in two separate experiments, MR-L2 reduced active lever presses and nicotine intake at both FR3 and FR5.

Conversely, chronic down-regulation of PDE4B protein, due to disruption of gene expression by induction of deletions and insertions in a coding region, resulted in the maintenance of nicotine IVSA (i.e., active presses and nicotine intake) despite high FR demand - which reduced IVSA in controls. We would expect chronic downregulation to have more complex effects on MSN function than acute administration of a drug such as MR-L2. cAMP signaling would be tonically enhanced, thus increasing nicotine and dopamine-dependent excitation of MSN. Additionally, resetting the transcriptional landscape and the downstream expression of neuronal proteins in multiple pathways would likely produce a myriad of changes in MSN function and in behavior that remain to be delineated. For example, *in vivo* transfection of constitutively active CREB has been shown to amplify the excitability of MSN (12).

The DaR subtypes (i.e. D1R and D2R) differentially expressed by two types of NAc MSNs (15–17) have opposite effects on cAMP synthesis (41). Activation of D1R induces adenylate cyclase to synthesize cAMP, whereas D2R inhibits this enzyme (41). Therefore, increased PDE4B activity would diminish D1R-dependent cAMP signaling and the activation of D1R-MSN; conversely, increased PDE4B activity would most likely amplify the inhibitory effect of dopamine on MSN that express D2R. Hence, PDE4B modulates both the activation and inhibition of MSN, depending in large part on the expression of D1R vs. D2R (15,16).

The effects of reducing the expression of PDE4B by CRISPR/Cas9 gene editing and enhancing the catabolic activity of the protein using MR-L2 depend on the subtype of MSN: a knock-down of the PDE4B gene would enhance D1R-dependent signaling and activation of D1R-MSN, whereas it would diminish the inhibitory effect of D2R signaling and reduce the inhibition of D2R-MSN by dopamine. Conversely, MR-L2 would be expected to reduce D1R-dependent signaling and increase D2R signaling. The relative role of each DaR-MSN subtype in determining the behavioral outcomes observed in this study are unknown. Both both subtypes of NAc MSN appear to be involved in reward and aversion (42–44), depending in part on the duration of stimulation (43). Moreover, the motivational effects of activating a specific subtype of DaR-MSN (i.e. D2R) depend on the phase of the motivated behavior (i.e. conditioned cue vs reward consumption) (45).

In summary, PDE4B plays a pivotal role in determining the NAcs motivational response to conditioned nicotine reinforcement. Reduced PDE4B activity, induced by CRISPR/Cas9 frameshift-induced gene inactivation, maintains the motivation to take nicotine as the cost of work increases, whereas increased PDE4B activity impairs the motivation to take the drug. Since NAc neuronal responses to goal directed behavior are heterogenous, depending on factors such as the specific neurons sampled, the operant paradigm, and the phase of the motivated behavior (31,32,45), the observations made in the present study may depend in part on the area of NAcs (i.e. dorsomedial) affected by CRISPR/Cas9 and MR-L2. Nonetheless, these reverse translational studies provide novel insights into the role of NAcs PDE4B in operant nicotine IVSA at increasing FR workload that contribute to understanding the heretofore unexplained GWAS associations between PDE4B and multiple stages of human smoking.

## Acknowledgments

Cas9+ female outbred Long Evans rats were generously provided by Professor Aron Geurts, Medical College of Wisconsin.

